## Supplementary figures and images for "Microvasculopathy, Luminal Calcification and Premature Aging in Fetuin-A Deficient Mice"

### Supplemental Figure1

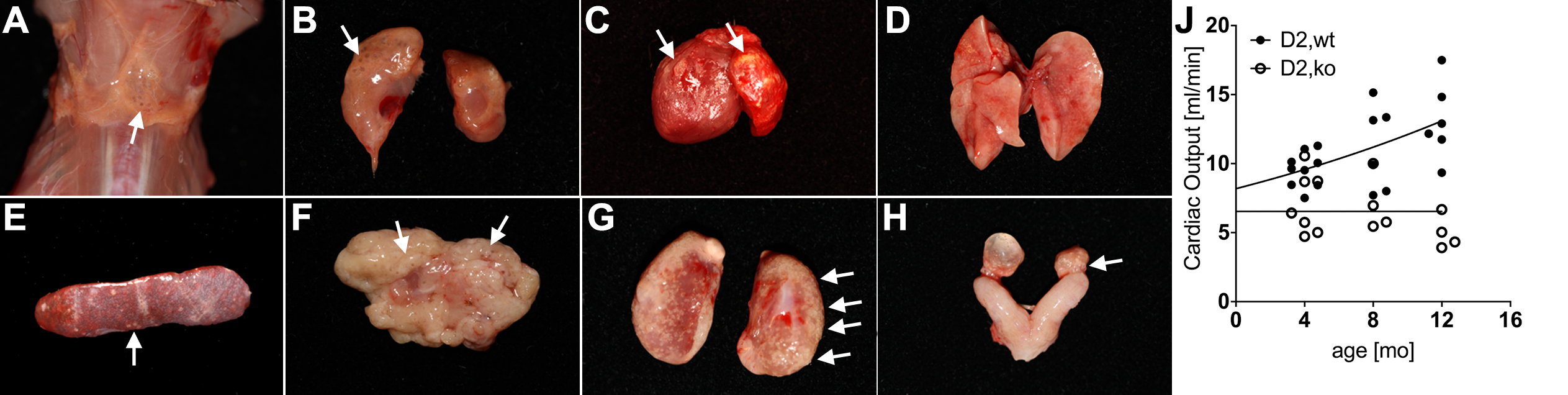

### Supplemental Figure2

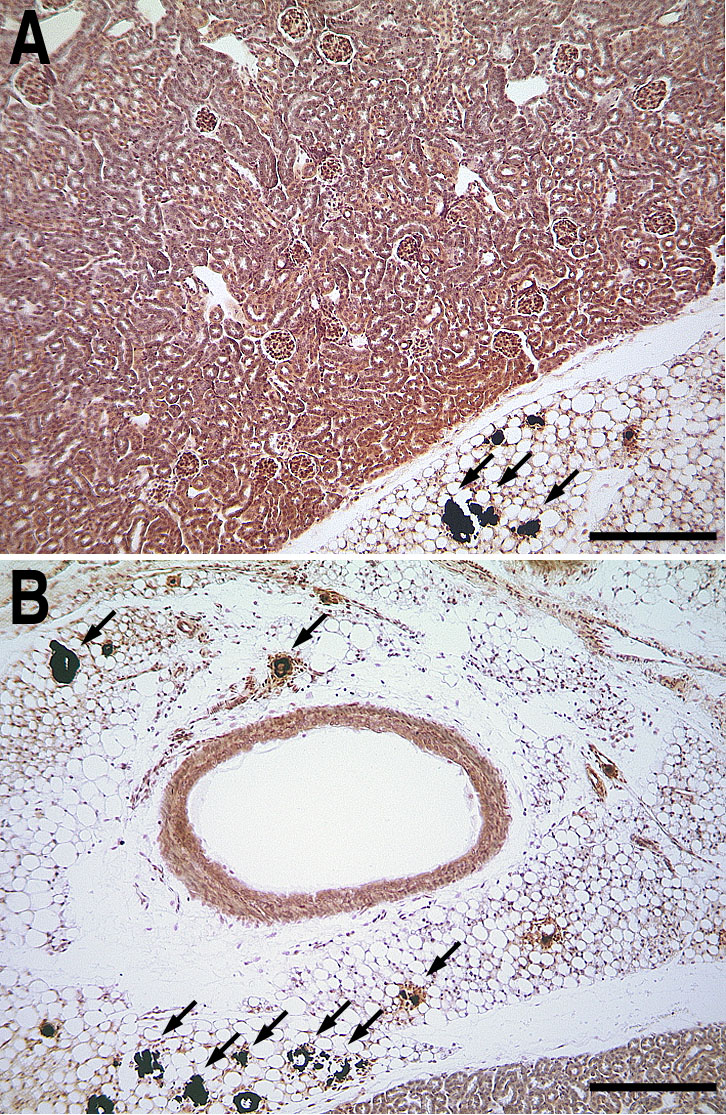

### Supplemental Figure3

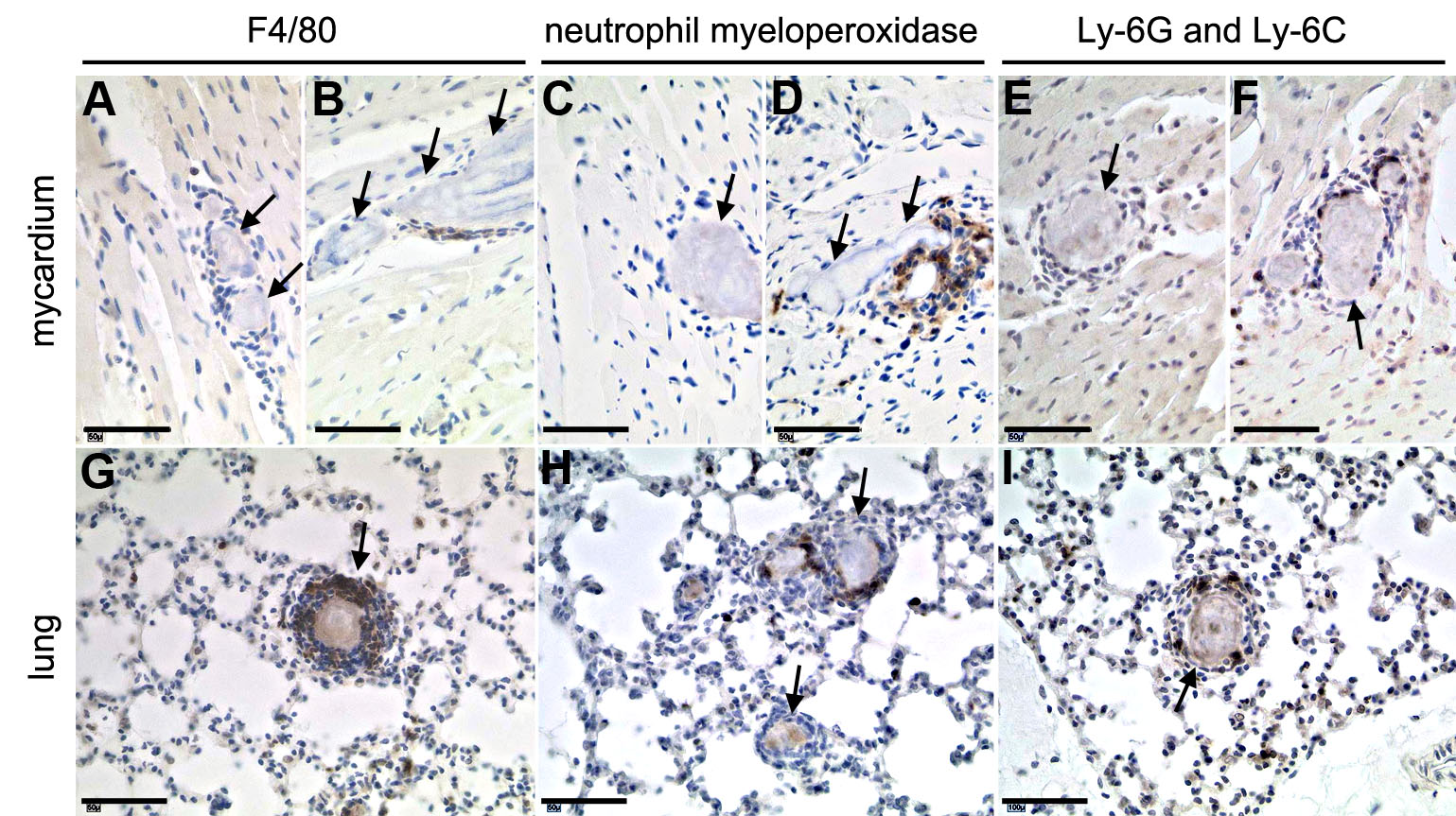
